# Estimation of Three-Dimensional Chromatin Morphology for Nuclear Classification and Characterisation

**DOI:** 10.1101/2020.07.29.226498

**Authors:** Priyanka Rana, Arcot Sowmya, Erik Meijering, Yang Song

## Abstract

Classification and characterisation of cellular morphological states are vital for understanding cell differentiation, development, proliferation and diverse pathological conditions. As the onset of morphological changes transpires following genetic alterations in the chromatin configuration inside the nucleus, the nuclear texture as one of the low-level properties if detected and quantified accurately has the potential to provide insights on nuclear organisation and enable early diagnosis and prognosis. This study presents a three dimensional (3D) nuclear texture description method for cell nucleus classification and variation measurement in chromatin patterns on the transition to another phenotypic state. The proposed approach includes third plane information using hyperplanes into the design of the Sorted Random Projections (SRP) texture feature. The significance of including third plane information for low-resolution volumetric images is investigated by comparing the performance of 3D texture descriptor with its respective pseudo 3D form that ignores the interslice intensity correlations. Following classification, changes in chromatin pattern are estimated by computing the ratio of heterochromatin and euchromatin corresponding to their respective intensities and image gradient obtained by 3D SRP. The proposed method is evaluated on two publicly available 3D image datasets of human fibroblast and human prostate cancer cell lines in two phenotypic states obtained from the public Statistics Online Computational Resource. Experimental results show that 3D SRP and 3D Local Binary Pattern provide better results than other utilised handcrafted descriptors and deep learning features extracted using a pre-trained model. The results also show the advantage of utilising 3D feature descriptor for classification over its corresponding pseudo version. In addition, the proposed method validates that as the cell passes to another phenotypic state, there is a change in intensity and aggregation of heterochromatin.

**Author Summary:** Automated classification and measurement of cellular phenotypic traits can significantly impact clinical decision making. Early detection of diseases requires an accurate description of low-level cellular features to detect small-scale abnormalities in the few abnormal cells in the tissue microenvironment. The challenge is the development of a computational approach for 3D textural feature description that can capture the heterogeneous information in multiple dimensions and characterise the cells in their ultimate and intermediate phenotypic states effectively. Our work has proposed the method and metrics to measure chromatin condensation pattern and classify the phenotypic states. Experimental evaluation on the 3D image set of human fibroblast and human prostate cancer cell collections validates the proposed method for the classification of cell states. Results also signify the credibility of proposed metrics to characterise the cellular phenotypic states and contributes to studies related to early diagnosis, prognosis and drug resistance.

## Introduction

Intercellular interactions within a tissue microenvironment take place through diverse mechanical and biochemical signals that control cellular development, differentiation and homeostasis [1]. Various studies have revealed that a derangement in signals disrupts chromatin dynamics and commences differential regulation of gene expression and genomic imbalance that subsequently triggers oncogenic transitions [1]. Chromatin domains on mutation undergo disarrangement of heterochromatin (HC) (condensed chromatin) organisation and observe coarsening or opening of HC, resulting in an increase or loss of HC aggregates all around the nuclear matrix region, respectively [2]. This transformation is closely associated with metastatic potential and holds diagnostic and prognostic significance. At the molecular level, alterations appear as a change in nuclear texture formed by wrinkles, folds and trenches manifested through entwined strands of nuclear proteins, lamins and chromatin. Changes in nuclear texture occur in conjunction with other morphological variations such as nuclear and nucleolar size, shape and count at the tissue level and the organisation of protein content, based on which cancerous cells are differentiated from normal ones [1, 2]. Although these alterations have been a gold standard for late-stage diagnosis of tumours, the description of their origin, interdependency and progression is not clear; therefore early diagnosis of cancer, drug discovery and prognosis care remains a challenge [1, 2].

The nucleus, being a prominent organelle of eukaryotic cells, houses cellular DNA (chromatin), hosts chromosome formation and offers a dynamic research domain to measure and study nuclear morphological changes from normal to malignant cells correlated with genetic alterations. The 4D Nucleome Network Project aims to create nuclear quantitative models that can provide insights into genome organisation in various cell types and their corresponding states in temporal and spatial domains [3]. In cross disciplines such as mechanobiology, nuclear morphological quantification has emerged as a promising approach to study the effect of external signals on nuclear morphology and their further impact on enclosed protein organisation. Quantitative analysis of variations in nuclear morphology and protein configuration helps to explain the mechanisms underlying cellular alterations and has opened new avenues for curative models in cancer care [1].

Quantitative models are usually evaluated on classification problems to comprehend and measure nuclear morphological alterations. Machine learning techniques that extract handcrafted features are often chosen over deep learning as the latter does not provide interpretable features [1]. Research on novel handcrafted feature descriptors thus remains active despite the success of deep learning [4–7], as they perform competently for well-defined problems and do not require massive amounts of data for training. Cellular changes at the molecular level can be understood through an adequate feature description capable of capturing low-level details of cells in multi-dimensions. The latest developments in confocal microscopy have enabled more effective in-vivo intraoperative studies through 3D fluorescence images, to analyse the heterogeneity of cellular patterns [2].

Nuclear texture, being one of the low-level properties when described and quantified accurately, has the potential to provide insights that enable early diagnosis and prognosis. Various studies have established the discriminative performance of 3D texture analysis more than 2D [8–10]. In the early years of 3D texture description, Gray-Level Co-Occurrence Matrix (GLCM) descriptors were widely used for fluorescence microscopy images [10]. Local Binary Pattern-Three Orthogonal Planes (LBP-TOP) [11] has been considered as the leading approach for 3D biomedical images [12]. It has outperformed commonly used, diverse variants of 3D texture descriptors such as Moments, Haralick, Tamura, Gabor and other variants of LBP for cell classification [5, 12, 13]. LBP-TOP computes standard LBP features from three orthogonal planes separately and concatenate them to represent 3D LBP feature descriptors. Apart from LBP-TOP, the recently proposed 3D RSurf [5] uses spherical coordinates with diverse combinations of azimuthal and polar angle to analyse 3D images. The latest scale and rotational invariant 3D Scale-Invariant Feature Transform (SIFT) [14] extracts texture features from a spherical image window, and is employed to target high intensity points of DAPI (4’,6-diamidino-2-phenylindole) images that represent HC.

In this study, we propose a novel approach to compute third plane information for texture description using cubic patches and hyperplanes within the patch. The proposed approach is adapted to the existing SRP feature descriptor [6] which describes the distribution of intensities in diverse patterns, originally defined for texture description of 2D images. SRP is based on the dimensionality-reduction technique known as Random Projections (RP), which compresses high dimensional data, captures salient information without information loss and preserves inter-distance of data values while projecting to a lower dimensional space. SRP is a rotationally invariant texture feature descriptor and has achieved competitive results and efficiency. The proposed 3D approach extends SRP by extracting features from the third plane using hyperplanes built in the cubic patches of the volumetric image. Bag of Visual Words (BOVW) [7] is then applied to generate the final 3D feature descriptor. The proposed 3D texture feature descriptor is used to measure changes in the chromatin patterns and perform cell nucleus classification, and evaluated on two publicly available 3D image datasets of DAPI stained nuclei of human fibroblast and human prostate cancer cells [15, 16]. Along with the earlier mentioned handcrafted features (SIFT, LBP and RSurf), Convolutional Neural Networks (CNNs) [17] are also used to generate deep learning features for comparison. The results demonstrate 3D SRP is one of the most effective texture feature description methods for this data set, and has potential to measure variations in chromatin patterns.

## Materials and Methods

### Data Description

In this study 3D Cell Nuclear Morphology Microscopy Imaging Dataset is used; the largest available public 3D image set obtained from Statistics Online Computational Resource (SOCR) [15, 16]. The dataset consists of two different cell collections, each comprised of different phenotypic states with distinct morphological features for binary classification (Fig 1). The first cell collection has 176 3D volumetric images of primary human fibroblast cells in two phenotypic states/classes: 1) 64 sub-volumes of proliferating fibroblasts (*PROLIF*), and 2) 112 sub-volumes of cell cycle synchronised by the serum-starvation protocol (*SS*). *SS* cells are obtained by subjecting fibroblasts cells to G0/G1 Serum Starvation Protocol which is known to produce changes in nuclear shape or size in human fibroblast cells [18]. The second cell line has 101 3D volumetric images of human prostate cancer cells (PC3) in two states/classes: 1) 50 sub-volumes of epithelial (*EPI*) state, and 2) 51 sub-volumes of mesenchymal transition (*EMT*) state. Both PC3 phenotypic states exhibit different and quantifiable nuclear morphological features that are useful in studying prostate cancer progression.

**Fig 1.**
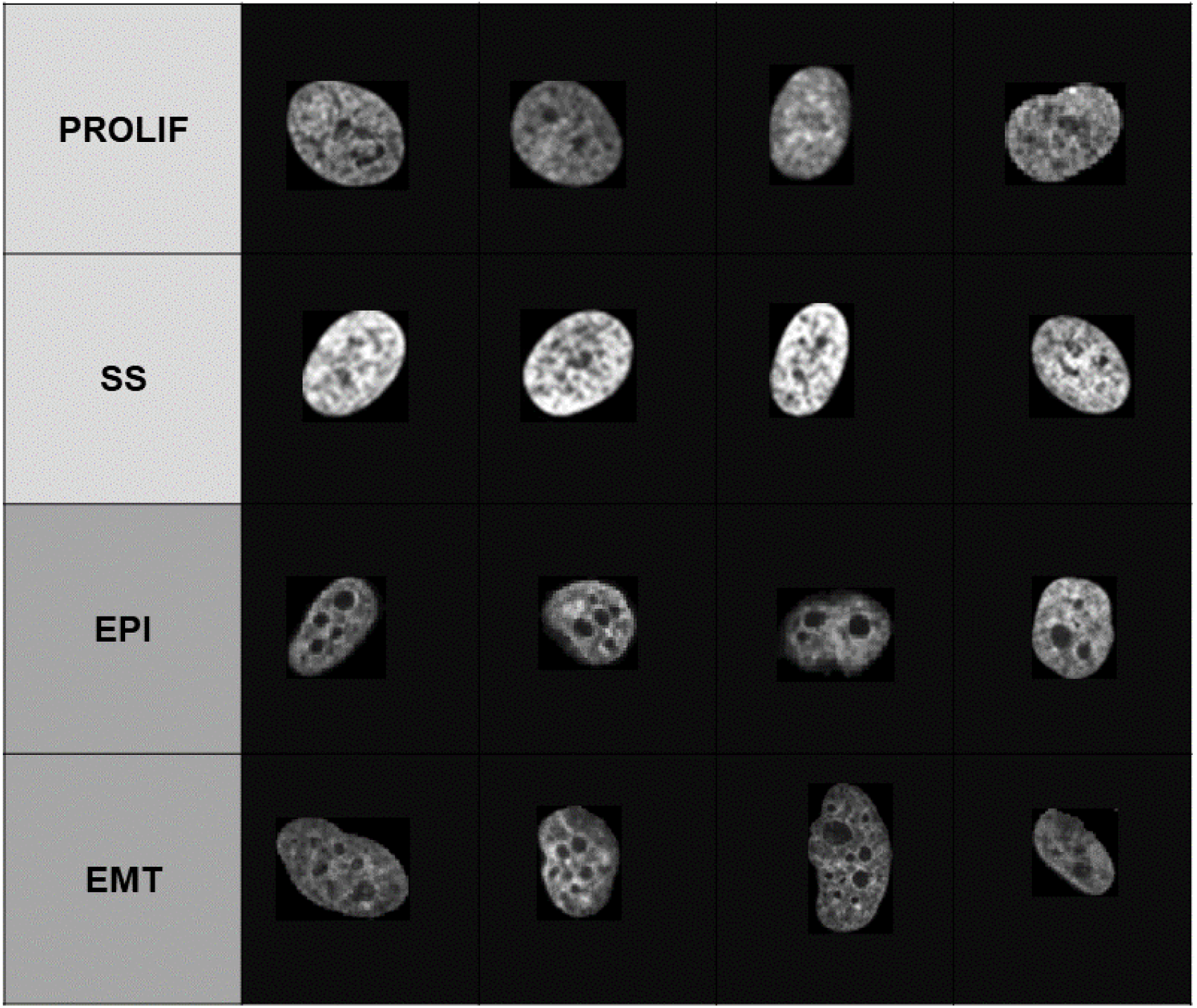
Representative sample images of Cell Nuclei.

All cell images have three channels showing different fluorophores: DAPI stain for nuclei, fibrillarin antibody (anti-fibrillarin) and ethidium bromide (EtBr) for staining nucleoli. Since the nucleolus contains very few pixels, we study only the DAPI images. Each volumetric image of both cell lines is in 3D TIFF format and has dimensions 1024 × 1024 × Z voxels, where Z ranges from 30 to 40 slices for fibroblast cell collection and 65 to 80 slices for PC3 cell collection. The voxel size of all volumetric images from both sets is 0.1318 × 0.1318 × 1 *µ*m^3^. The dataset also includes meta-data extracted from the original data.

### Data Preprocessing and Segmentation

As a first step, segmentation of individual cell nuclei from potentially noisy microscopic images is performed before classification. At times, uneven staining may cause inaccurate segmentation, therefore as a quality control protocol, nuclei at the border of the image or without nucleoli are excluded. Slices from the volumetric image are extracted using ImageJ [19] and fed into CellProfiler [20]. The CellProfiler modules (Fig 2A) identify the nuclei and nucleoli and relate them, followed by filtering out of unwanted nuclei. Considering uneven illumination in the images, adaptive thresholding with Otsu’s method [21] is employed. The approach classifies image pixels into three classes, namely background, foreground and middle intensity, where the middle intensity has the option to be merged into the foreground or background in the final segmentation result. The CellProfiler output window shows corresponding results that are promising for closely positioned objects (Fig 2B). In addition, instances of connected nuclei are resolved by applying the watershed algorithm [22] wherever required (Fig 2C).

**Fig 2.**
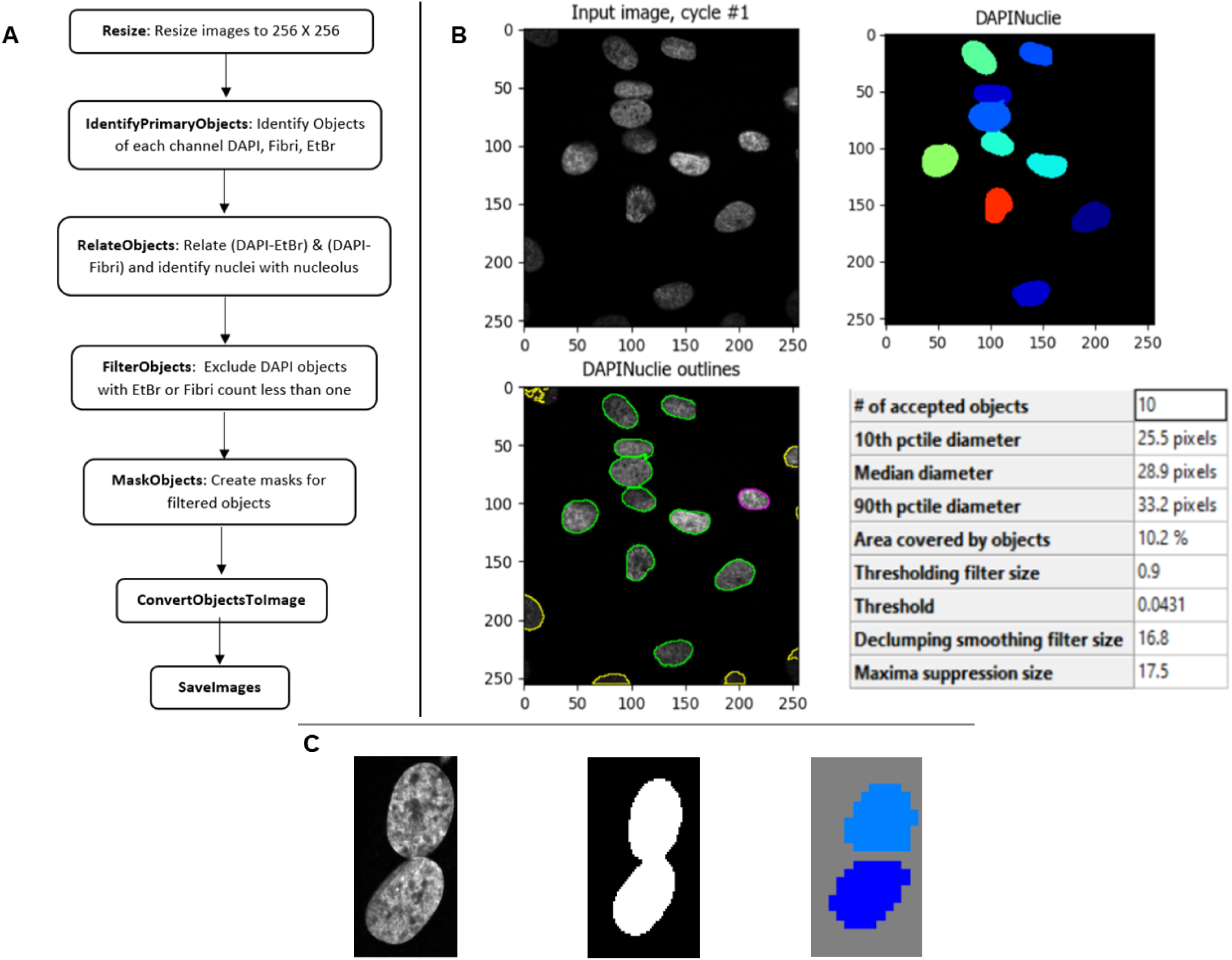
Segmentation. A: Framework of CellProfiler modules. B: CellProfiler output window for segmentation. C: Example Image; corresponding mask; segmentation by watershed algorithm.

Following segmentation, cell objects are cropped from the image slices and stacked to form a 3D cell object for classification. The average dimension of cropped cell objects across all samples is 64 × 64 pixels. Due to parameter setting for the Otsu algorithm, objects with very low intensity could not be identified in a few top-most and bottom-most slices, resulting in final object thickness up to 15 slices for fibroblast cells and 60 for PC3 cells.

A comparison of the achieved segmentation results with previous studies [15, 16] on the same dataset is shown in Table 1. In the current results, 22 more *SS* cell objects, 25 fewer *PROLIF* cell objects, 103 and 141 more *EMT* and *EPI* cell objects are detected, respectively. According to Kalinin et al. [15], their segmentation results do not represent ground truth, as they are not hand labelled by an expert. Their study discarded connected cell objects; however, this study employed a semi-automated approach where results from CellProfiler are visually inspected, and the watershed algorithm is applied to segment connected cell objects, leading to a higher number of SS, *EMT* and *EPI* cell objects. CellProfiler, being a highly sensitive tool, has discarded comparatively more *PROLIF* cell objects, based on applied quality protocols.

**Table 1.**
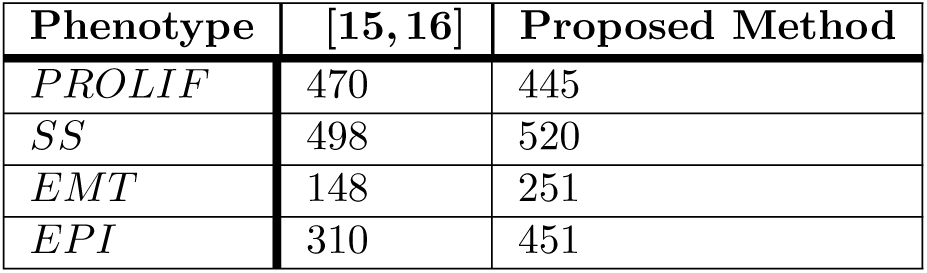
Segmentation Results.

### Feature Description

In this study, handcrafted feature descriptors (SRP, LBP, SIFT and RSurf) and transfer learning to obtain deep learning features are used to extract texture features from the data. This section elaborates the proposed approach of 3D extension of the SRP feature descriptor. The other descriptors and feature representation techniques are also briefly discussed.

### SRP and 3D SRP

SRP is a rotationally invariant texture feature descriptor, which describes the distribution of intensities in diverse patterns. It is defined by five functions (Fig 3B) computed by accessing image pixels in a global, circular and square pattern, and pixel differences in an angular and radial pattern. Values from each ring are sorted, concatenated and projected to a lower dimensional space while preserving the original distances between data points [6]. Dimensionality reduction is performed via RP which projects points from a high-dimensional space to a randomly created lower dimensional space using the *RP* matrix and a linear operation.

**Fig 3.**
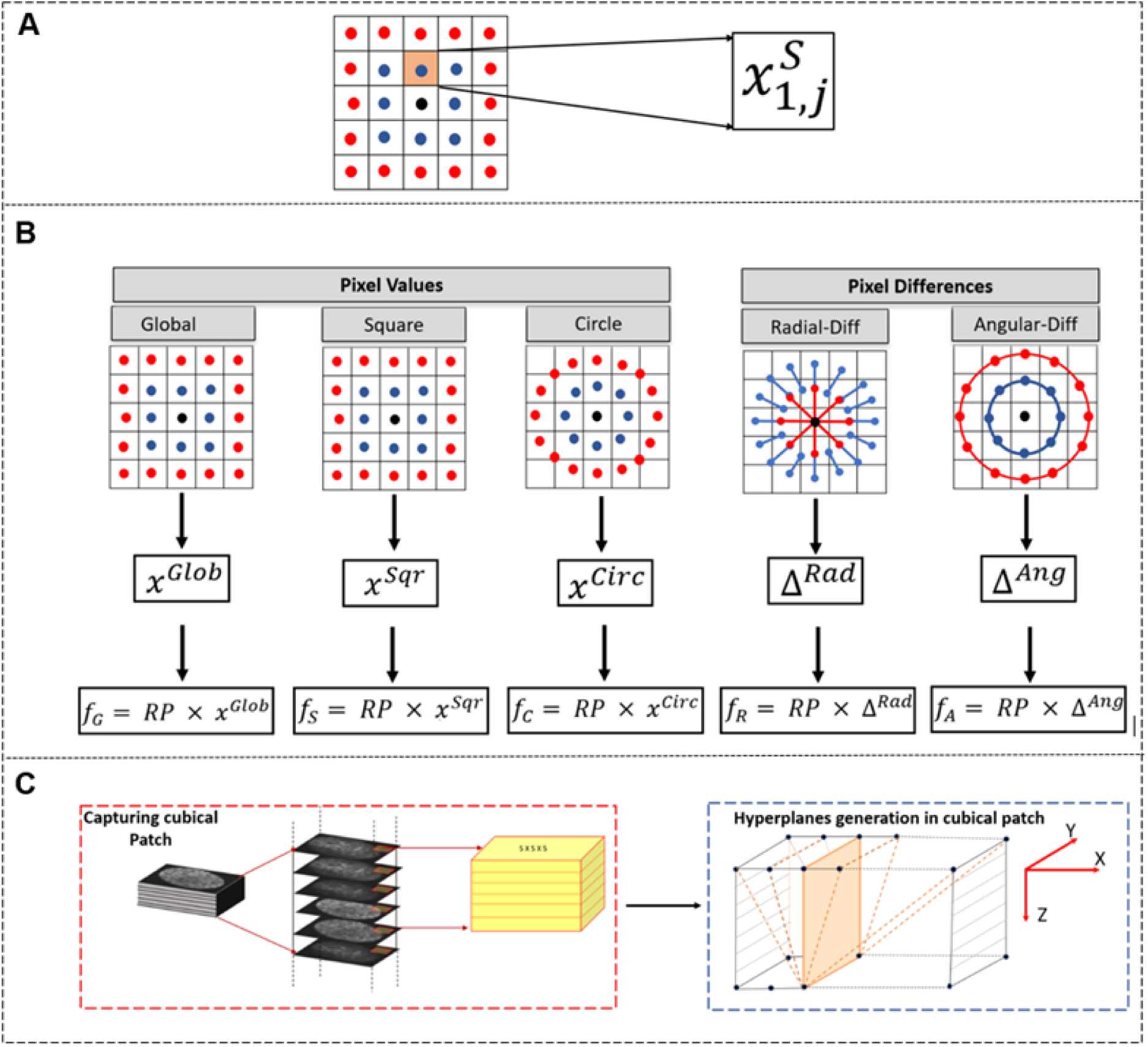
SRP. A: Represents pixel value in 1^*st*^ Square ring (*S*) at *j*^*th*^ position. B: Example image patch of 5 x 5 pixels; local sorted descriptors computed using Eq (3); computation of SRP feature functions. C: Proposed 3D SRP approach.

RP is designed based on the theory of compressed sensing, which states that any non-adaptive linear measurement in the form of random projections is capable of preserving intrinsic and salient information of the original compressible signal in lower dimensional space [6]. *RP* matrices can be constructed in various ways. In this work, the method employed follows Medeiros et al. [23] where the *RP* matrix elements (*rp*)_*ij*_ are obtained randomly as

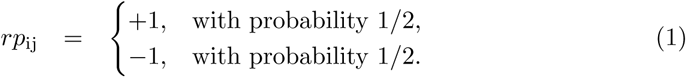

Accordingly, the high dimensional data vector *D* of dimension *b* is transformed to lower dimensional space *L* utilising the *RP* matrix with dimensions *a* × *b* and simple linear operation :

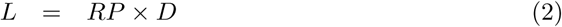

where the resultant dimension of *L* is *a*. The value of *a* in the dimension of RP is correlated with the patch size of the image. As SRP features are extracted in concentric circles and squares (Fig 3B), options for patch size dimensions are odd numbers as [(2*n*+1) × (2*n*+1)] where *n* = 2,3, … *N* and depend on the original dimensions of the input image and the level of the desired description. The value of *a* equals 10 for *n*=2 and for every increment in *n, a* increases by 10. These are the optimal RP dimensions for corresponding patch sizes and have been empirically confirmed and presented by Liu et al. [24] under different normalisation techniques.

As illustrated in Fig 3A, 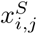 is the pixel in the *i*th square ring (*S*) in *j*th position, *x*^*Sqr*^ is the final vector built by concatenating sorted pixels of *m* concentric squares and *p*_*i*_ is the number of pixels in the *i*th square. This is represented as

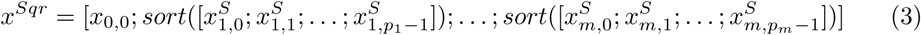

Similarly, *x*^*Circ*^, Δ^*Ang*^ and Δ^*Rad*^ are computed for pixels in circular pattern, angular pixel difference, and radial pixel difference, respectively. *x*^*Glob*^ is obtained by sorting all the pixels of the patch. Accordingly, the feature vector of a square pattern *f*_*S*_ is computed through a linear operation:

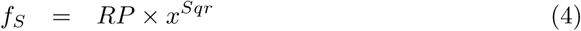

Similarly, *f*_*G*_, *f*_*C*_, *f*_*A*_ and *f*_*R*_ are computed for global, circular, angular and radial patterns, respectively. The angular and radial differences between pixels preserve the inter-distance of neighbouring pixels, making SRP feature description invariant to rotations.

Unlike other texture descriptors utilized in this study, there is currently no 3D version of SRP. To extract 3D SRP features, the existing 2D SRP is thus extended and the third plane information is proposed to be accessed using cubic patches and building multiple hyperplanes within the patch. For 3D SRP, as shown in Fig 3C, a cubic patch is extracted from a 3D volumetric image with dimensions 5 × 5 × *z*, where *z* in this implementation is 5, the same value as the dimensions in *X* and *Y* planes. For images with slices fewer than [2(2*n*+1)], *z* can be the same as the number of slices. As demonstrated in Fig 3C, multiple hyperplanes are generated between each column of the bottom-most slice and columns of the top-most slice in a cubic patch. Pixel coordinates between the top and bottom slices are identified using 3D Bresenham’s line algorithm [25]. Five SRP functions for each hyperplane 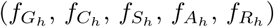 are computed and concatenated across all hyperplanes built along the *Y Z* plane of the cubic patch.

The resultant vectors 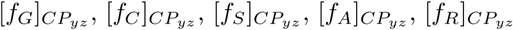 (*CP* stands for cubic patch) are each normalised into 16 bin histograms and combined to represent the SRP descriptor of the *Y Z* plane of a cubic patch with dimension 80 (16 bins × 5 functions):

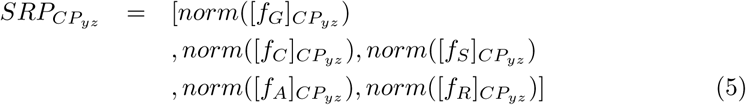

This process is repeated for the *XZ* plane as well. Features from the *XY* plane are extracted from the slices following the same computational sequence. Subsequently, descriptors from the three planes are concatenated to obtain the final 3D SRP feature descriptor for a cubic patch with dimension 240(80 bins x 3 planes):

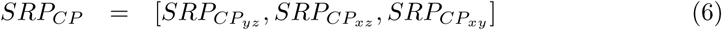

The patch-based feature description results in a large set of local feature descriptors of cubic patches which are often redundant and could deteriorate the classification performance. Since SRP features are generated from the RP matrix, they are random and already globalised. However, their coupling with non-local statistical feature descriptors such as BOVW can make them robust and improve their classification performance. Hence following BOVW, feature vectors of cubic patches from all volumetric images are clustered to generate a visual vocabulary of the most informative and dominant features by utilising *k*-means and sum pooling (details in the Feature Representation section). Subsequently, each 3D image is represented by the visual vocabulary word that is identified based on the closest distance between the image feature and visual words in the codebook. In this way, the volumetric image represented by a set of cubic patches is embedded into a compact vector space.

### Other Texture Descriptors

This study includes a comparison of the proposed approach with other handcrafted feature descriptors, which employ distinct approaches for third plane inclusion and are comparatively recent, such as RSurf, and the widely utilised LBP and SIFT texture descriptors. The choice of LBP and SIFT is also motivated from a recent review of texture descriptors that mentions SIFT and LBP as milestone texture feature descriptors [7]. In order to achieve a reasonable comparison, the same patch size (5 × 5 pixels) as 3D SRP is used for 3D LBP. RSurf features are extracted from the whole image.

LBP features for this study are computed following LBP-TOP [11], a 3D version of LBP which suggests computation of standard LBP features from three orthogonal planes separately and concatenate them to represent 3D LBP feature descriptors. Following Zhu et al. [26], neighbourhood and radius of 16 and 2 are used respectively for 5 × 5 patch size.

Computation of the 3D SIFT [14] feature vector utilises a spherical image window of radius 2*σ* (the constant multiple of different scales of the scale space pyramid) and its centre as the keypoint. To represent image data, the gradient histogram is generated for each cubical sub-region (dimensions 4 × 4 × 4 voxels) of the spherical image window. The gradient histogram for each subregion is computed with 12 directions per histogram; therefore, the dimension of the final feature vector is (4 x 4 x 4) x 12 = 768. The utilised dimensions of the cubical sub-region and histogram length in this study have been defined as the optimal parameters in the original work, which detects the maximum number of keypoints.

RSurf [4] features, one of the latest additions to the list of handcrafted features, captures intensity variations by traversing the image in different directions. The traversed pixel vector is represented by four functions (length of the vector, difference between the highest and lowest intensities, sum of all intensity values, and the number of times the intensity varied from high to low and vice versa). As shown in Fig 4, for pseudo 3D RSurf, the implementation in this work considered four angles (0*°*, 45*°*, 90*°*, 135*°*) to traverse the intensities in each slice. For non-pseudo 3D RSurf [5], spherical coordinates where azimuthal angle varies from 0*°* to 360*°* at intervals of 45*°* and polar angle from -90*°* to 90*°* at intervals of 30*°* have been used. Diverse values of angles enable rotational invariance.

**Fig 4.**
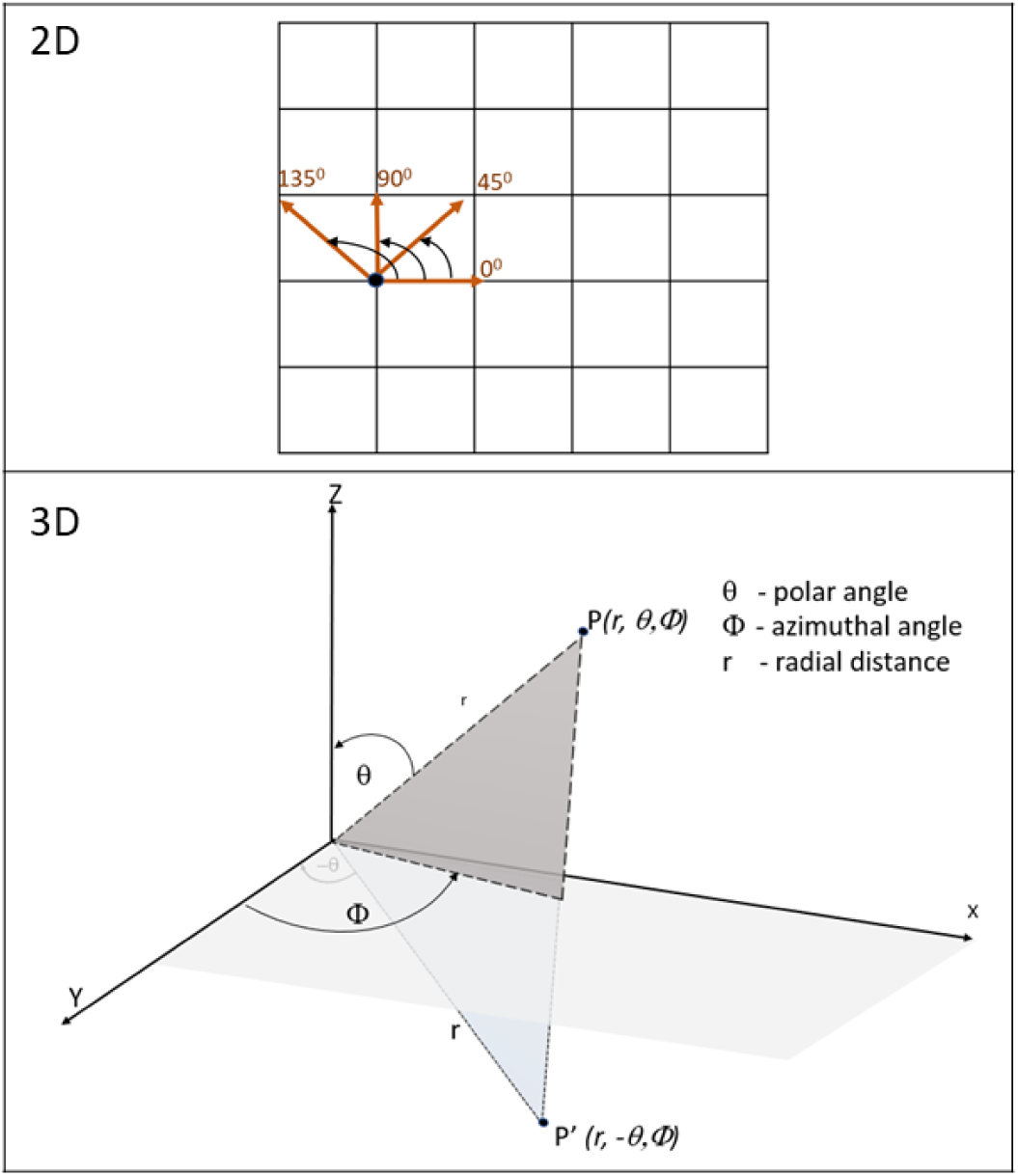
RSurf. Traversal of pixels in 2D and 3D..

**Fig 5.**
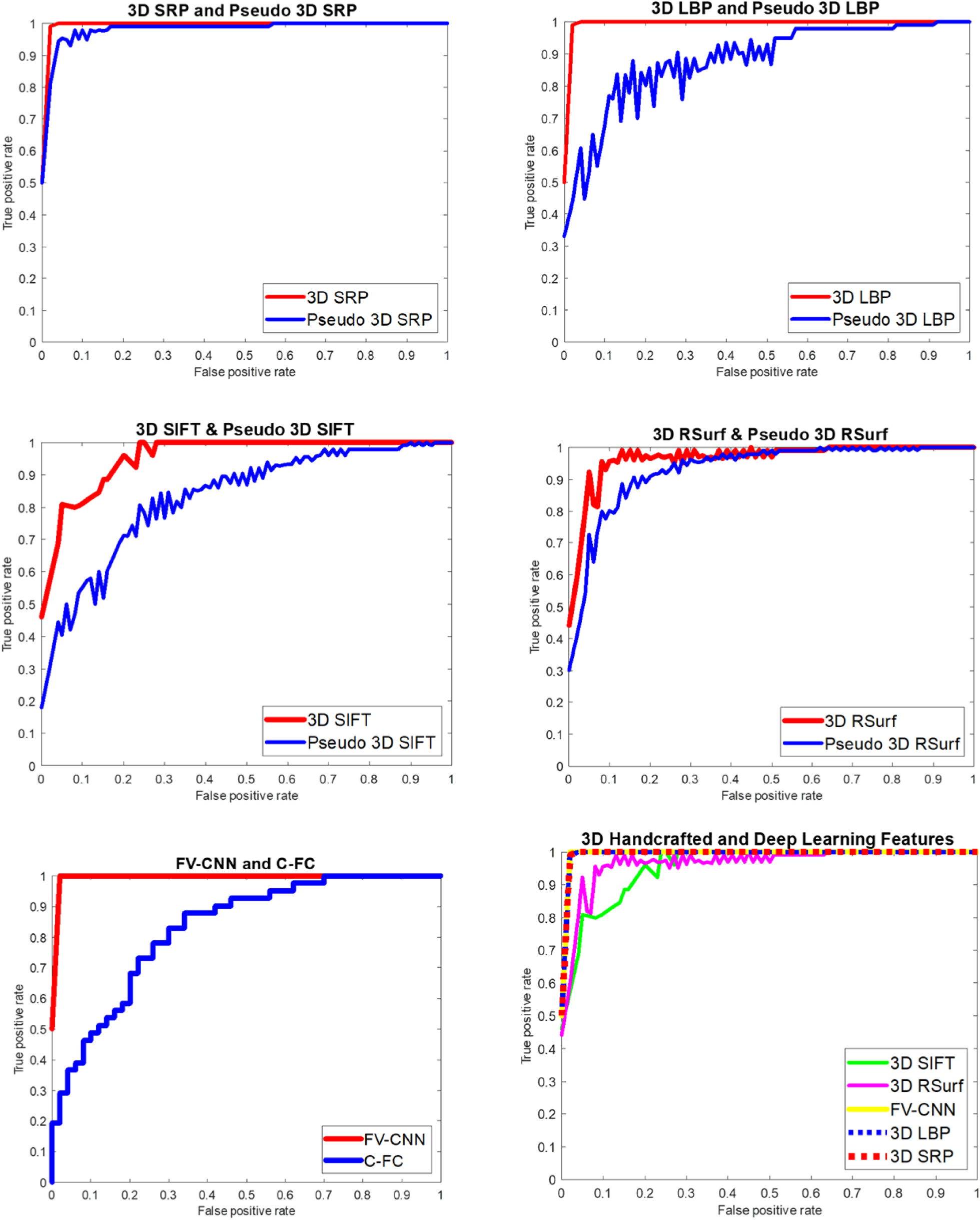
(A) ROC Curves. Fibroblast Dataset.

**Fig 5.**
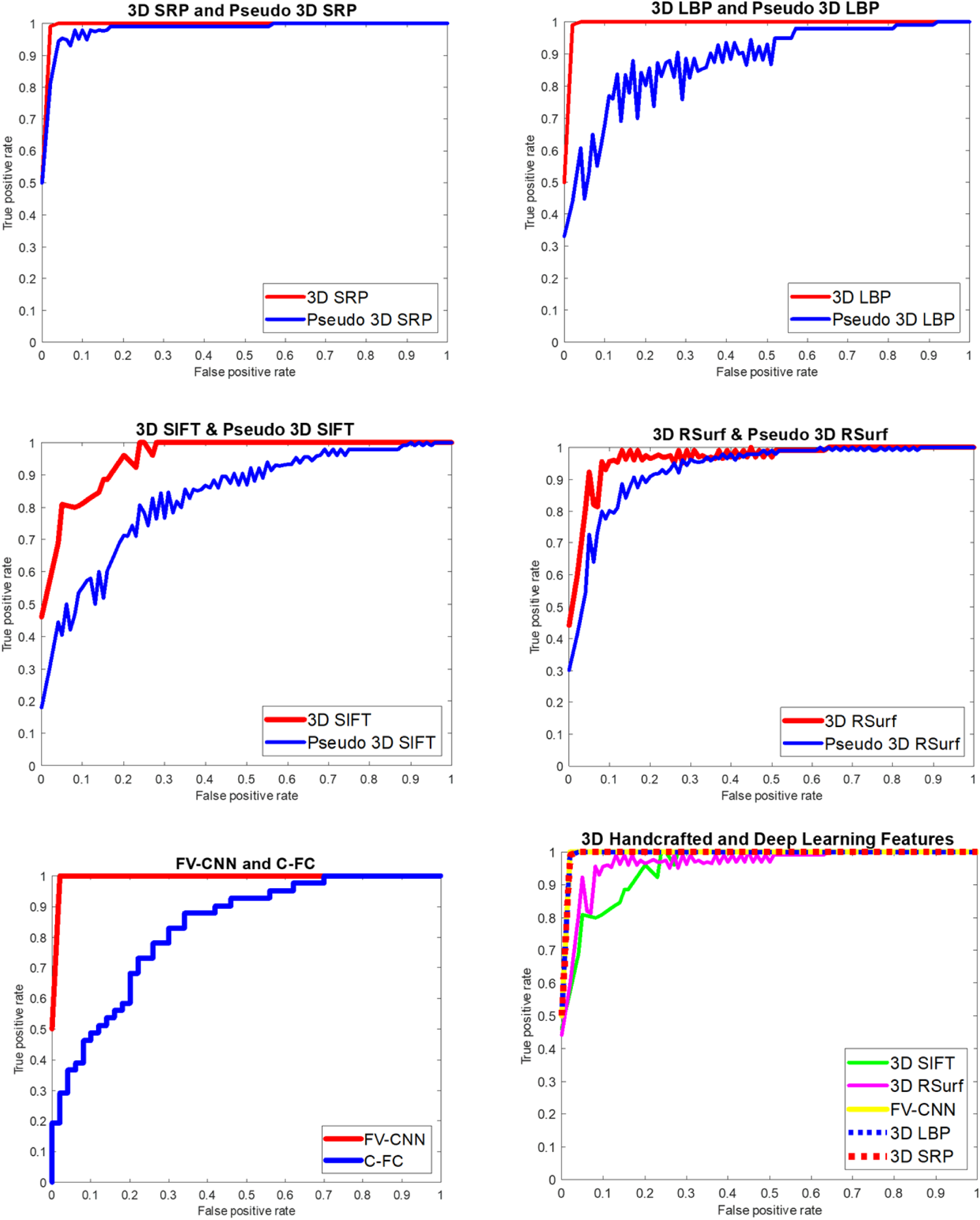
(B) ROC Curves. PC3 Dataset.

### Deep Learning

In this study, deep learning features are also evaluated by applying the transfer learning technique. Transfer learning reduces the complexity of obtaining deep learning features and has shown success in various biomedical studies either by fine-tuning the network on available data or as a feature extractor. VGG-16, a CNN model trained on ImageNet, has established itself as a reliable choice in biomedical studies [27]. Using it directly for feature extraction does not demand any changes to the architecture by the user and provides robust deep learning features [28, 29]. With VGG-16, the input image goes through a series of convolutional layers before it finally produces a dense set of local feature descriptors of 512 dimensions at the last fully convolutional layer. In this study, 3D features are generated with the pre-trained VGG-16 model by encoding local features of last convolutional layer from all slices of the 3D image using the Fisher Vector (FV) [30, 31] which produces the resultant CNN based FV descriptor (FV-CNN) with dimension 65536 (details in the Feature Representation section). Classification results based on the features obtained from the penultimate fully-connected layer (C-FC) of the CNN model are also included for comparison.

This study is focused on texture description of fluorescent images, therefore, FV-CNN features have been included only to compare the proposed approach with deep learning features. While the usual practice to obtain CNN features in biomedical studies is by training a new CNN model, the available number of images poses a limitation on training an optimal CNN. Since this study is not focussed on CNN design, experiments with a pre-trained CNN model are conducted instead.

### Feature Representation

To overcome high intraclass and low interclass variations, robust texture description is required, alongside compact and efficient representation to make it useful in real life applications. Feature extraction from image patches often results in a large number of features, with many being redundant, leading to high computational cost and poor discriminative strength. Therefore feature representation techniques such as BOVW [7] and FV [30, 31] encoding are employed which group local feature descriptors into elements of a codebook that encodes many similar local features into high-level features.

BOVW is one of the popular techniques for effective and efficient feature representation to find the most informative features. The current implementation utilises *k*-means clustering for codebook generation, an unsupervised learning algorithm to generate the visual vocabulary as clusters, and sum pooling to quantise the image in the form of a histogram vector. Before codebook generation, an appropriate value of *k* is obtained following the elbow method. In this method, *k*-means clustering is performed and the Sum of Squared Error (SSE) is obtained for different values of *k* (32, 64, 96, 128, 160). A graph of SSE for each value of k is plotted, which usually takes the shape of an arm, and the value of k corresponding to the elbow of the arm is chosen as an optimal value which represents the least value of *k* after which SSE scarcely varies. BOVW encoding is applied to the pseudo and non-pseudo versions of SRP, LBP, SIFT and RSurf and the resultant feature dimension is 64.

The FV descriptor encoding employs Gaussian Mixture Models (GMM) [29], a probability density function for codebook generation. Unlike BOVW where the image is represented by the number of occurrences of the visual word, FV encodes the gradient of the log likelihood of features with respect to the GMM parameters (mean vector, standard deviation vector and mixing weights). The current implementation generates G = 64 Gaussian components from patch-level features. The final FV descriptor of each image is the concatenation of the derivatives with respect to GMM parameters, resulting in the dimension of the FV encoding of each image as 2GD, where D is the dimension of the local feature vector. This FV encoding is applied to the deep learning (CNN) features, and the resultant dimension of FV-CNN is 65536 (2 x 64 x 512). Considering the high dimension of the FV encoding, we used BOVW for handcrafted features. FV for deep learning features is employed following the work of Song et al. [29].

### Classification

Cellular alterations at the molecular level may happen inconsistently in a small group of defective cells in a tissue microenvironment. Therefore, image classification based on cellular feature descriptors is recommended to comprehend the variations. Support Vector Machine (SVM) classification model is trained on the various texture feature descriptors, utilising the Radial Basis Function (RBF) as the kernel function for automatic cell nucleus classification. *SS* and *EMT* are labelled as the positive group; *PROLIF* and *EPI* represent the negative group. In order to evaluate the performance of cellular textural feature descriptors for volumetric image-based classification, 10-fold cross-validation is used, with all cells from one 3D image included in the same fold. There are 45-50 cells per fold for fibroblasts cell collection and 25-45 cells per fold for PC3 cell collection.

Classification performance is evaluated using AUC (Area-Under-the-Curve) of the ROC (Receiver-Operating-Characteristic) curve and the F1 score. ROC is plotted for TPR (True Positive Rate) vs FPR (False Positive Rate) to demonstrate classifier performance at different threshold settings, and AUC measures the area under the curve. Evaluation of overall effectiveness of the classification model is also measured by the Recall (percentage of positives correctly identified by the classifier) and Precision (percentage of true positives out of the identified positives) metrics. The F1 score considers false positives and false negatives and balances Precision and Recall by computing their harmonic mean:

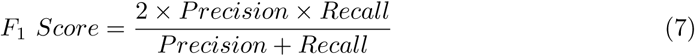

### Measurement of Changes in Chromatin Pattern

Chromatin patterns vary according to cell cycle phase, developmental state and chromatin positioning [32]. HC is characterised by the constriction in centromere, defined by a variant of histone H3, centromere protein-A (CENP-A) which is an established biomarker [33]. Being a profoundly condensed and compact chromatin fraction, it is easily detectable by DAPI staining [34, 35]. Thus corresponding to the high luminance contrast regions in DAPI images, HC is identified by applying a threshold equal to the sum of the minimum intensity and sixty per cent of the difference between the maximum and minimum intensities [36].

Following the proposed 3D SRP, this study takes two approaches based on pixel values and adjacent pixel differences, to determine the chromatin pattern alterations in 3D. The maximum value of the SRP feature descriptor functions computed from each 5 × 5 × 5 cubic patch is employed to estimate the threshold. The threshold value is used to identify HC, followed by computing the ratio of HC to EC corresponding to the respective pixel values and pixel differences obtained from SRP functions. The patch size can be determined according to the respective classification performance. HC/EC ratio of both cell states is statistically compared to inspect the nature of changes in chromatin pattern on transition to another phenotypic state.

### HC Intensity Measure

HC/EC ratio based on pixel values estimates the fraction of the HC intensity in the nucleus. Following Eq (4), for an image with *n* cubic patches, the global SRP descriptor for the *i*^*th*^ patch is computed as 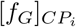, and the corresponding maximum value is obtained as *m*_*i*_. The threshold to identify intensity corresponding to HC is taken as the least *m*_*i*_ of all the respective values of the cubic patches of the volumetric image. There are cells oriented at a certain angle which makes *m*_*i*_ for corner patches equal to zero, hence the threshold is set as the minimum non-zero value. The global feature descriptor vectors obtained from *n* cubic patches of the nuclear image [*f*_*G*_]_*Nucleus*_ are traversed, and the values above the threshold are identified corresponding to HC ([*f*_*G*_]_*HC*_). Subsequently, the fraction of the heterochromatin intensity is estimated using the following equation:

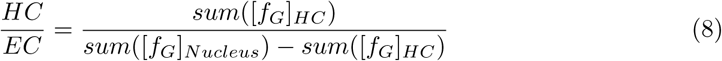

### HC Aggregation Measure

HC/EC ratio based on pixel differences estimates the fraction of the HC aggregates in the nucleus. The increase or decrease in image gradient leads to the corresponding rise or drop in the slope of the curve of HC aggregate, which in turn indicates condensation or decondensation of chromatin, respectively. As described in the SRP section, the radial and angular descriptor function for the *i*^*th*^ patch of an image with n cubic patches is computed as 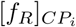 and 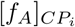, respectively. Following this, the maximum value of the concatenated vector 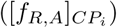, is obtained as *m*_*i*_. Similar to the intensity measure, the threshold to identify image gradient corresponding to HC is taken as the least *m*_*i*_ of all the respective values of the cubic patches of the volumetric image. The radial and angular feature descriptor vectors obtained from *n* cubic patches of the nuclear image ([*f*_*R,A*_]_*Nucleus*_) are traversed, and the values above the threshold are identified corresponding to HC ([*f*_*R,A*_]_*HC*_). Subsequently, the fraction of the HC aggregates is estimated using the following equation:

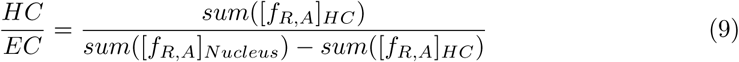

## Results

The results shown in Table 2 are the mean and standard deviation of the metric values measured on the classification based on 10-fold cross-validation. In Table 3, the best achieved results are shown in comparison to the state-of-the-art (SOTA) [15, 16]. While the current work evaluates texture descriptors using 10-fold cross-validation, the compared SOTA works performed 20 split Leave-2-Opposite-Groups-Out (L2OGO) cross-validation and performed classification with 968 fibroblast cell objects (498 *SS* and 470 *PROLIF*) and 458 PC3 cell objects (148 *EMT* and 310 *EPI*). Due to the different approach adopted in this work for segmentation and quality protocols, 965 fibroblast cell objects (520 *SS* and 445 *PROLIF*) and 702 PC3 cell objects (251 *EMT* and 451 *EPI*) are obtained for classification. The number of cells in each subsample for the current study is 45-50 for fibroblast cells and 25-45 for PC3 cells, while Kalinin et al. [16] used 19 cell sets. The SOTA works also merged features from all the channels by quantifying nucleoli-level features for each nucleus using statistical values such as average, median, maximum and higher moments, while the current study is focussed only on the DAPI channel, as the number of pixels from other channels is too small to be considered for texture description.

**Table 2.** SVM Classification Results for Fibroblasts and PC3 Cell Collection.

|  | FIBROBLAST CELL LINE |  |  |  | PC3 CELL LINE |  |  |  |
| --- | --- | --- | --- | --- | --- | --- | --- | --- |
| Features | AUC | F1 Score | Recall | Precision | AUC | F1 Score | Precision | Recall |
| *Pseudo 3D SRP* | 0.94±0.04 | 0.94±0.04 | 0.97±0.04 | 0.91±0.06 | 0.98±0.01 | 0.85±0.20 | 0.75±0.32 | 0.94±0.08 |
| *3DSRP* | **1±0** | **0.99±0.02** | **0.98±0.03** | **1±0** | **1±0** | **0.98±0.02** | **0.97±0** | **0.98±0.03** |
| *Pseudo 3DLBP* | 0.88±0.13 | 0.79±0.15 | 0.77±0.18 | 0.82±0.10 | 0.97±0.02 | 0.90±0.06 | 0.84±0.11 | 0.97±0.04 |
| *3DLBP* | **1±0** | **0.99±0.02** | **0.98±0.03** | **1±0** | 0.99±0 | 0.97±0.03 | 0.97±0.04 | 0.97±0.04 |
| *Pseudo 3D SIFT* | 0.85±0.04 | 0.74±0.08 | 0.69±0.10 | 0.79±0.08 | 0.93±0.02 | 0.80±0.13 | 0.84±0.21 | 0.80±0.05 |
| *3D SIFT* | 0.96±0.02 | 0.90±0.05 | 0.90±0.06 | 0.91±0.13 | 0.98±0.03 | 0.96±0.06 | 0.97±0.04 | 0.94±0.07 |
| *Pseudo 3D RSurf* | 0.94±0.02 | 0.86±0.02 | 0.87±0.05 | 0.85±0.05 | 0.97±0.03 | 0.89±0.05 | 0.89±0.01 | 0.91±0.08 |
| *3D RSurf* | 0.98±0.02 | 0.92±0.03 | 0.93±0.04 | 0.92±0.02 | 0.97±0.03 | 0.88±0.09 | 0.85±0.12 | 0.93±0.09 |
| *FV – CNN* | **1±0** | 0.93±0.16 | 0.89±0.21 | 1±0.01 | **1±0.01** | 0.96±0.06 | 0.95±0.03 | 0.96±0.08 |
| *C – FC* | 0.72±0.08 | 0.62±0.08 | 0.61±0.11 | 0.65±0.08 | 0.76±0.05 | 0.72±0.04 | 0.70±0.06 | 0.76±0.08 |

**Table 3.**
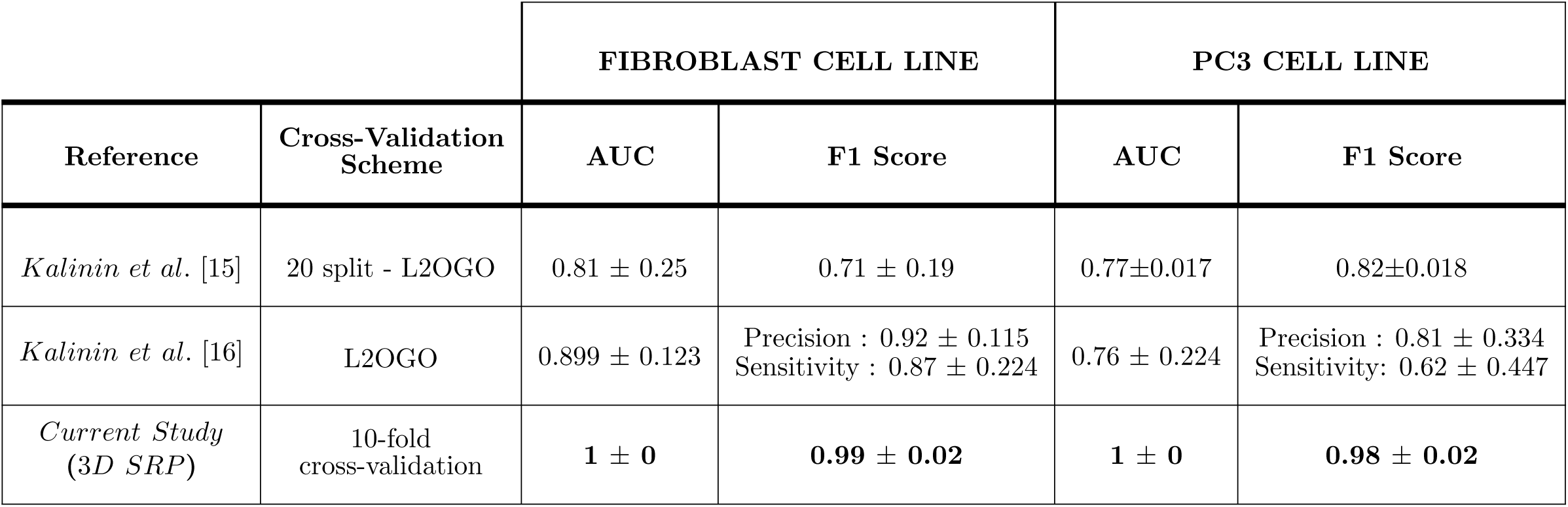
Comparison of classification results with existing studies.

|  |  | FIBROBLAST CELL LINE |  | PC3 CELL LINE |  |
| --- | --- | --- | --- | --- | --- |
| Reference | Cross-Validation Scheme | AUC | F1 Score | AUC | F1 Score |
| *Kalinin et al.* [15] | 20 split - L2OGO | $0.81 \pm 0.25$ | $0.71 \pm 0.19$ | $0.77 \pm 0.017$ | $0.82 \pm 0.018$ |
| *Kalinin et al.* [16] | L2OGO | $0.899 \pm 0.123$ | Precision : $0.92 \pm 0.115$<br>Sensitivity : $0.87 \pm 0.224$ | $0.76 \pm 0.224$ | Precision : $0.81 \pm 0.334$<br>Sensitivity: $0.62 \pm 0.447$ |
| *Current Study*<br>(3D SRP) | 10-fold cross-validation | **$1 \pm 0$** | **$0.99 \pm 0.02$** | **$1 \pm 0$** | **$0.98 \pm 0.02$** |

Variability in performance differences between pseudo and non-pseudo versions of the feature descriptor is demonstrated for all the folds through the ROC plots in Fig 5(A) and Fig 5(B). Classifiers for all the non-pseudo versions of texture feature descriptors performed better than their corresponding pseudo versions, except pseudo RSurf which is as good as non-pseudo 3D RSurf for the PC3 data set. Since slice count for PC3 cells is up to 60, the number of considered angular values for the polar angle is not sufficient to extract all the pixel information. As a result, several pixels in the top slices are left out when the angle is projected from the bottom most slices, and vice versa. Therefore, objects with a large slice thickness require more diverse polar angles with smaller intervals for effective feature description, and this may lead to high computational cost. Based on the AUC and F1 score values, non-pseudo 3D LBP and 3D SRP performed better than all other considered hand-crafted feature descriptors on both data sets. In comparison, FV-CNN features show an overlapping ROC curve with 3D SRP and 3D LBP and demonstrate distinctly higher performance than C-FC.

### Statistical Evaluation of Classification Results

The two-sample t-test at 1% significance level is used to compare AUC and F1 scores of pseudo and non-pseudo descriptors based on the null hypothesis:

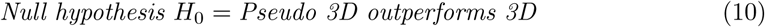

As shown in Table 4 and Table 5, the test results verify the statistical significance of the advantage of non-pseudo versions of descriptors over the pseudo versions successfully except RSurf on the PC3 image set.

**Table 4.**
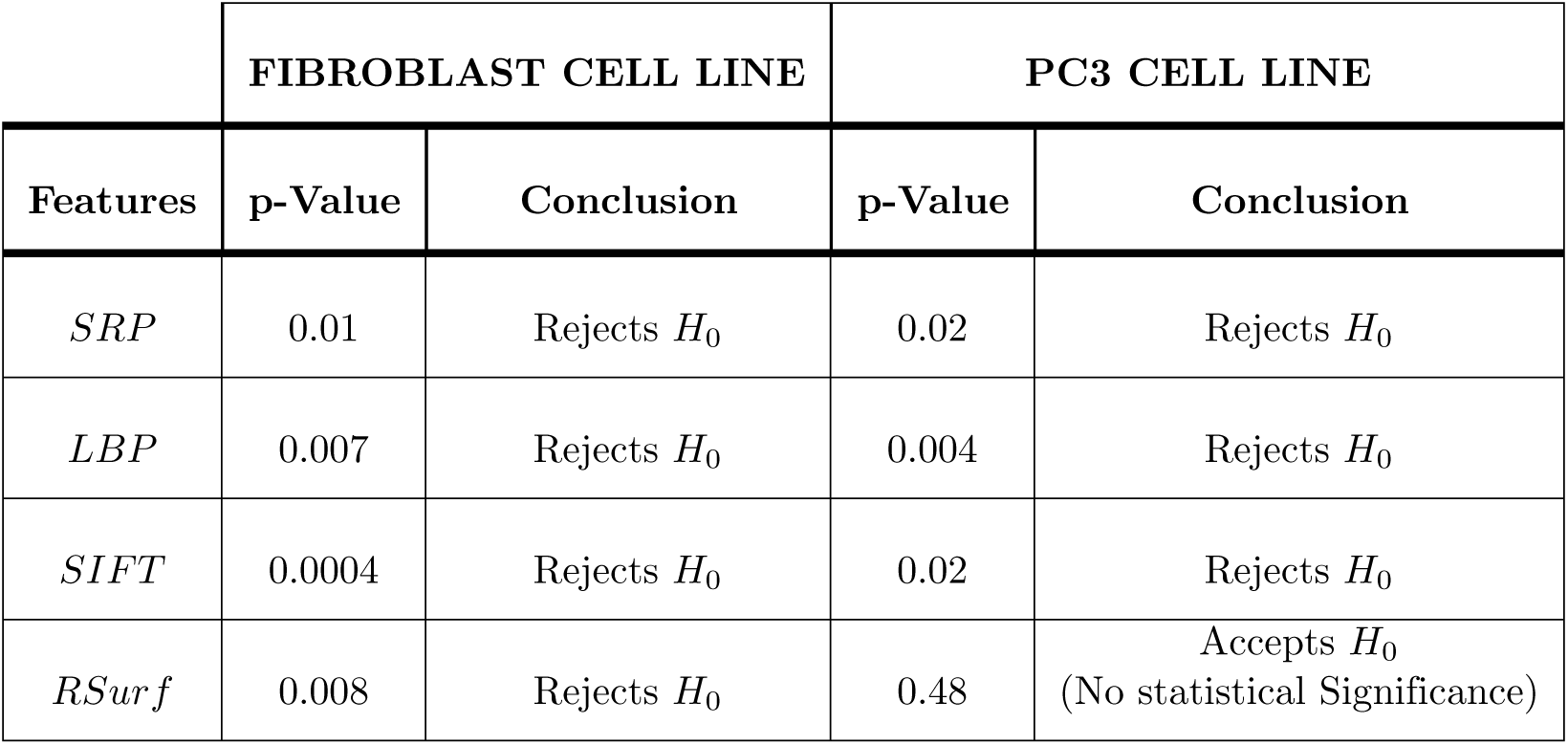
Two sample t-test results for AUC.

**Table 5.** Two sample t-test results for F1 Score.

|  | FIBROBLAST CELL LINE |  | PC3 CELL LINE |  |
| --- | --- | --- | --- | --- |
| Features | p-Value | Conclusion | p-Value | Conclusion |
| *SRP* | 0.002 | Rejects $H_0$ | 0.02 | Rejects $H_0$ |
| *LBP* | 0.002 | Rejects $H_0$ | 0.004 | Rejects $H_0$ |
| *SIFT* | 0.001 | Rejects $H_0$ | 0.005 | Rejects $H_0$ |
| *RSurf* | 0.004 | Rejects $H_0$ | 0.69 | Accepts $H_0$<br>(No statistical Significance) |

Since 3D versions of feature descriptors outperformed their corresponding pseudo forms, inter distinctiveness of non-pseudo 3D descriptors is explored in further analysis. As not all samples satisfy the assumption of homoscedasticity, the Kruskal-Wallis test [37] is employed, which is a non-parametric method also called one-way ANOVA based on ranks, to compare the means of more than two sample groups. The hypothesis for the Kruskal-Wallis test is:

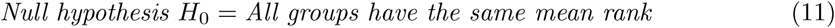

At 1% significance level and very small p-value (< 0.01), the Kruskal-Wallis test indicates the difference between mean ranks of five groups (SRP, LBP, SIFT, RSurf, FV-CNN). In Fig 6 and Fig 7, different colours of the lines indicate different mean ranks and population. The length of the line represents the comparison interval, and the extent of overlap of lines implies the range of similarity of corresponding groups.

**Fig 6.**
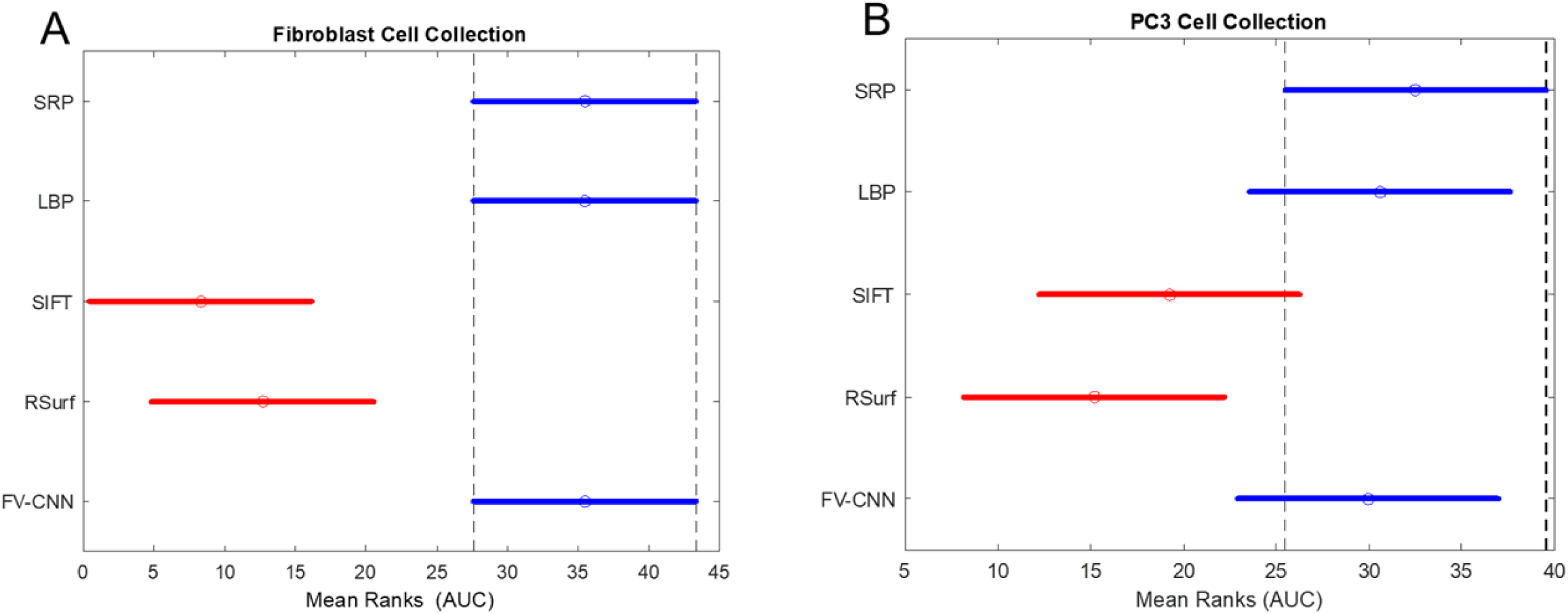
Mean ranks of 3D handcrafted descriptors and FV-CNN from Kruskal – Wallis test results for AUC. A: Fibroblast Cell Collection. B: PC3 Cell Collection.

**Fig 7.**
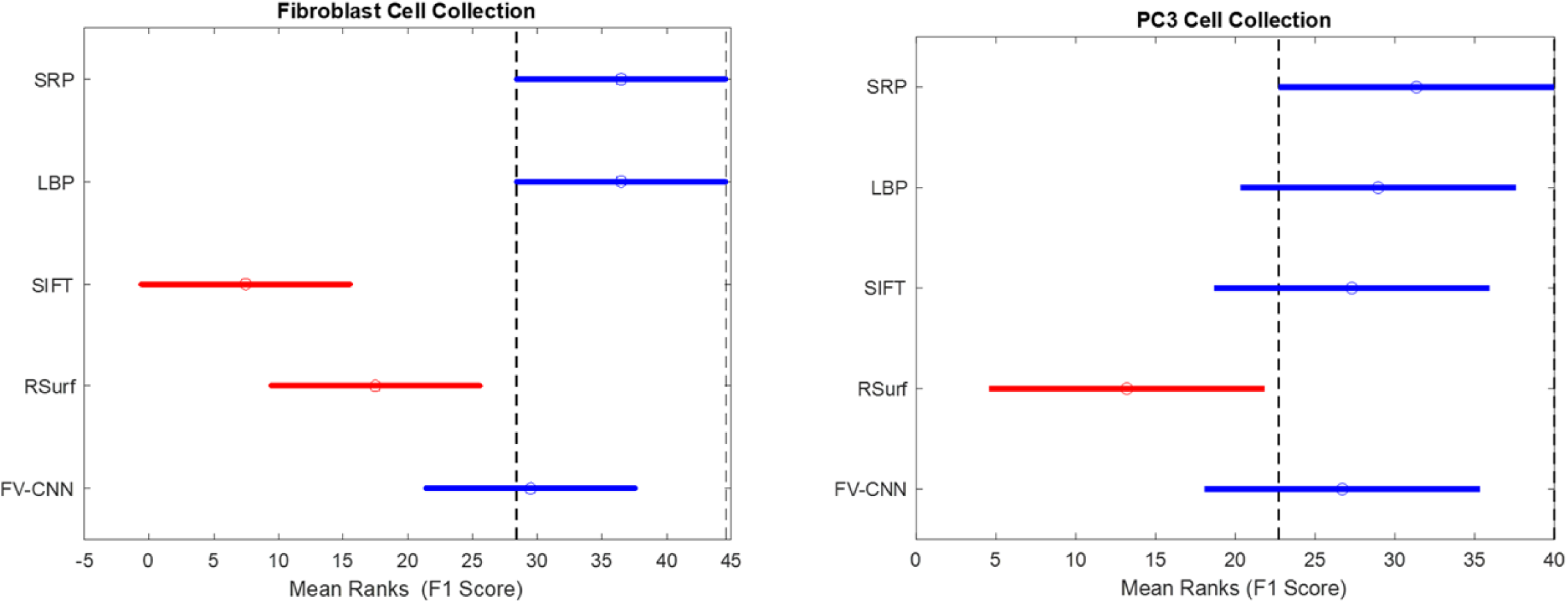
Mean ranks of 3D handcrafted descriptors and FV-CNN from Kruskal – Wallis test results for F1 Score. A: Fibroblast Cell Collection. B: PC3 Cell Collection.

As demonstrated in Fig 6A and Fig 6B, 3D LBP, 3D SRP and FV-CNN are at similar ranks and from a separate population than 3D RSurf and 3D SIFT. Although the mean F1 score for FV-CNN is lower than 3D SRP and 3D LBP for fibroblast cell images, statistical results demonstrate their performance to be similar (Fig 7A). This is because the F1 score of two subsamples deteriorated the mean FV-CNN. However, all other subsamples performed as well as 3D SRP and 3D LBP. F1 scores for most of the feature descriptors are good for the PC3 data set with a high number of slices (Fig 7B). Overall 3D SRP and 3D LBP performed better than all other considered handcrafted feature descriptors, while 3D SRP achieved better results than 3D LBP for the PC3 data set.

### Statistical Evaluation of HC/EC

The two-sided Wilcoxon rank-sum test at the significance level of 1% is utilised to measure the difference in *HC/EC*_*P ixelV alues*_ and *HC/EC*_*PixelDifferences*_ between two classes for both cell lines. The corresponding null hypothesis is:

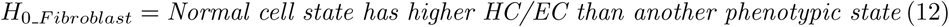

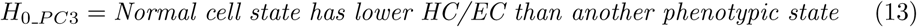

As shown in Table 6, the results from the test verify the statistical significance of the difference in *HC/EC*_*PixelV alues*_ and *HC/EC*_*PixelDifferences*_ between two classes and indicate both ratios are higher for *SS* cells than for *PROLIF* (normal state) cells, while they are lower for *EMT* cells than for *EPI* (normal state) cells. The utilised data set has multiple subsets of images for each class where each subset represents one ‘run’ of the microscope of different cell samples. The test was conducted for all individual subsets, and the observations remained the same for all subsets except in one instance when the test was performed excluding one image subset (coded as 179) comprising 25 volumetric images with 134 cells in the *EMT* class of PC3 data set. *PC*3^179^ refers to the image set which includes only this subset and PC3* refers to the image set without this subset in EMT class. On evaluation for PC3* image set with 117 cells, we observed high p-values which indicates insufficient evidence to reject the null hypothesis for *HC/EC*_*P ixelV alues*_ and *HC/EC*_*P ixelDifferences*_. However, the *z*-value (value of the *z*-statistic) of 0.6341 indicates a negative shift in the median of *HC/EC*_*P ixelDifferences*_ from *EPI* to *EMT* class at 1% significance level.

**Table 6.**
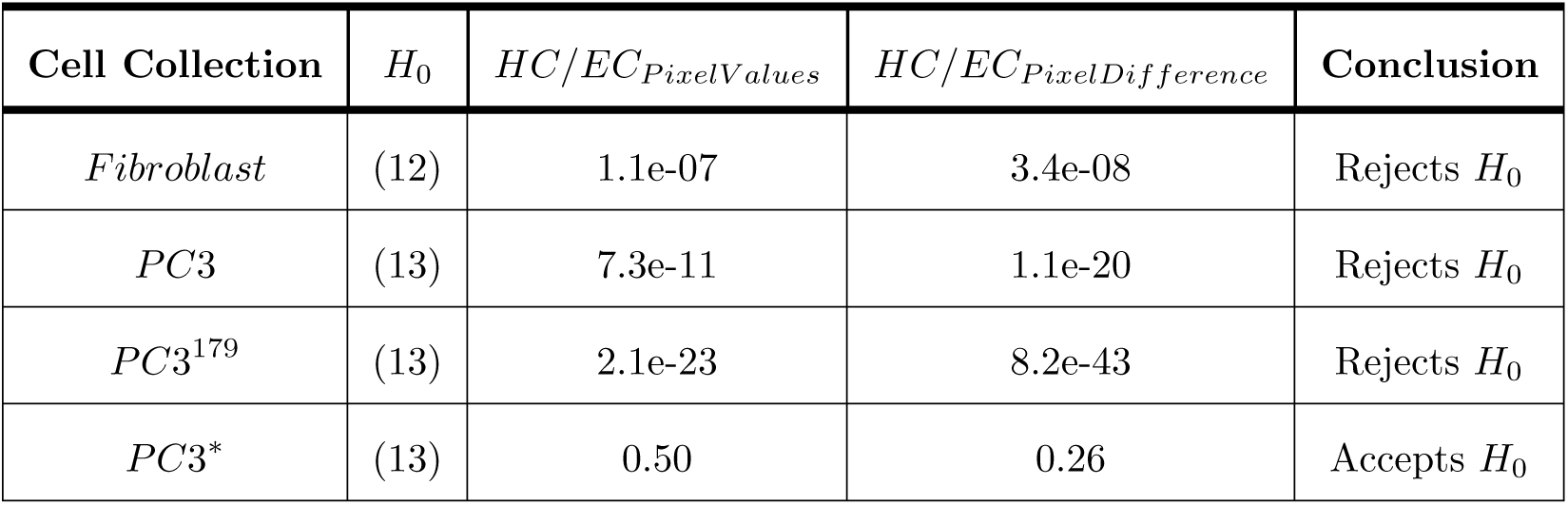
Two sample t-test results for F1 Score.

The *HC/EC* ratio was also computed following the same method with the original pixel values without random projections, and it was observed that the obtained p-value was higher. However, the inference remains the same for the pixel differences approach and no statistical significance in ratio difference is found for the pixel value method. *HC/EC* ratios were also computed following the pseudo 3D approach. Similar to the proposed 3D method, the obtained results demonstrated a positive shift in *HC/EC*_*P ixelV alues*_ and *HC/EC*_*P ixelDifferences*_ from *PROLIF* to *SS* class and negative shift from *EPI* to *EMT* class. However, resultant p-values were higher than the values obtained from the proposed 3D approach, suggesting the latter is more credible.

## Discussion

In this study, 3D handcrafted features along with deep learning features were utilised for the classification of phenotypes. 3D versions of SIFT, LBP, RSurf and SRP were compared with their corresponding pseudo forms to study their effectiveness in 3D and infer the impact of computing third plane information for low resolution. To extract information from the third plane, RSurf utilises spherical coordinates, SIFT utilises spherical image window, LBP combines standard LBP features from all three planes, while the proposed approach applied to SRP extracts cubic patches from the volumetric image and builds multiple hyperplanes within the patch. Experimental results demonstrate the advantage of utilising 3D feature descriptors for classification over their corresponding pseudo versions. Although the 3D image data has low resolution along the *Z* plane, results show that it still contributes to improved results. It is noticed that feature descriptors based on traversing the intensity of the image in patches perform better than non-patch-based image descriptors such as RSurf or SIFT. On comparing the performance of a feature descriptor on two data sets (fibroblast and PC3 cell collections), it is observed that the approach to include third plane information and the corresponding input parameters are significantly correlated with the number of slices in the volumetric image and must be carefully considered to obtain the best classification performance. For example, 3D SIFT based on spherical image window exhibited better performance for PC3 data set images with a higher number of slices, as the slices were adequate to build an appropriate sphere. In contrast, 3D Rsurf’s performance deteriorated, as the considered number of polar angles were not sufficient to extract all the pixel information. Similarly, 3D SRP outperformed 3D LBP on the PC3 data set, as its capability to extract extensive information was fully utilised for the images with a higher number of slices. In addition, SRP features offer more robust feature description as these are obtained by encoding intensity distributions of an image patch in square, circular and inter ring structure, while LBP utilises only pixel differences in circular pattern.

Obtained metric values include AUC = 1 ± 0, F1 score = 0.99 ± 0.02, precision = 1 ± 0, recall = 0.98 ± 0.03 for fibroblasts and AUC = 1 ± 0, F1 score = 0.98 ± 0.02, precision = 0.97 ± 0 and recall = 0.98 ± 0.03 for PC3 cell lines, which are higher than the current SOTA (fibroblast data set : AUC = 0.899 ± 0.123, precision = 0.922 ± 0.115, recall (sensitivity) = 0.874 ± 0.224; PC3 data set : AUC = 0.899 ± 0.123, precision = 0.922 ± 0.115, recall = 0.874 ± 0.224) obtained by classification using shape-based features on the same dataset. Factors such as quality protocols in the preprocessing step, the segmentation approach and feature representation primarily contribute to the high values of AUC and F1 score. Image background and noise often degrade the power of texture descriptors, therefore feature extraction in this study has been performed on cell objects instead of whole image. The implementation of quality protocols using CellProfiler, which is an advanced tool for microscopic images, eradicates noise and segments closely positioned objects effectively. The adopted segmentation approach is a semi-automated, two-step process where the second step applies the Watershed algorithm to connected cell objects identified on visual inspection of the Cell Profiler output images. Redundancy in features significantly slows down the classifier training, and it is addressed using feature encoding techniques such as BOVW and Fisher Vectors.

Understanding of nuclear topology and the organisation of subnuclear components motivated the development of the hypothesis that changes in shape and volume of the nucleus for the considered data [15, 16] would demonstrate considerable changes in nuclear texture too. The results shown in Table 3, establish the hypothesis, and prove that texture features are stronger features than shape and size for the classification of nuclear morphology. Nuclear morphology is supported by structural proteins called lamins that lie underneath the inner nuclear membrane and chromatin the enclosed genetic material [1]. As evident from earlier studies [15, 16], *SS* imposes significant changes in cell size and shape, which refer to changes in lamins, the protein primarily responsible for nuclear size and shape. The presented experiments demonstrate changes in intrinsic texture mainly formed by chromatin; this indicates the molecular linkage of lamins and chromatin proteins and the correlation of the possible textural changes with shape and size alterations. Since *EMT* happens in an early stage of cancer development, our study also addresses the role of texture descriptors in early cancer diagnosis. Results for the PC3 data set based on texture descriptors demonstrate higher discriminative strength than previous studies based on shape and size, and this emphasises the prominent potential of texture descriptors to catch molecular level variations in early diagnosis and prognosis.

Notably, low p-values obtained from statistical analysis of HC/EC ratios indicate the significant increase in intensity and HC aggregation, signifying HC condensation on the transition to *SS* state (Fig 8(A)). For the PC3 data set, on the other hand, there is a decrease in the intensity and HC aggregation, which signifies a decondensation or “open chromatin” state on the transition to the *EMT* state (Fig 8(B)). Results in Table 6 imply changes in the chromatin pattern, which is in accordance with other relevant studies [1, 2, 8, 36]. It was also observed that the p-value was higher when HC/EC ratios were computed based on their original intensity values instead of values in the random projections. This is because RP achieves compact representation of high dimensional data and preserves its inter-distances when projecting values to a lower dimensional space. This intensifies the salient information of the data, providing global features, and therefore, statistical analysis demonstrated the distinction of ratios more significantly.

**Fig 8.**
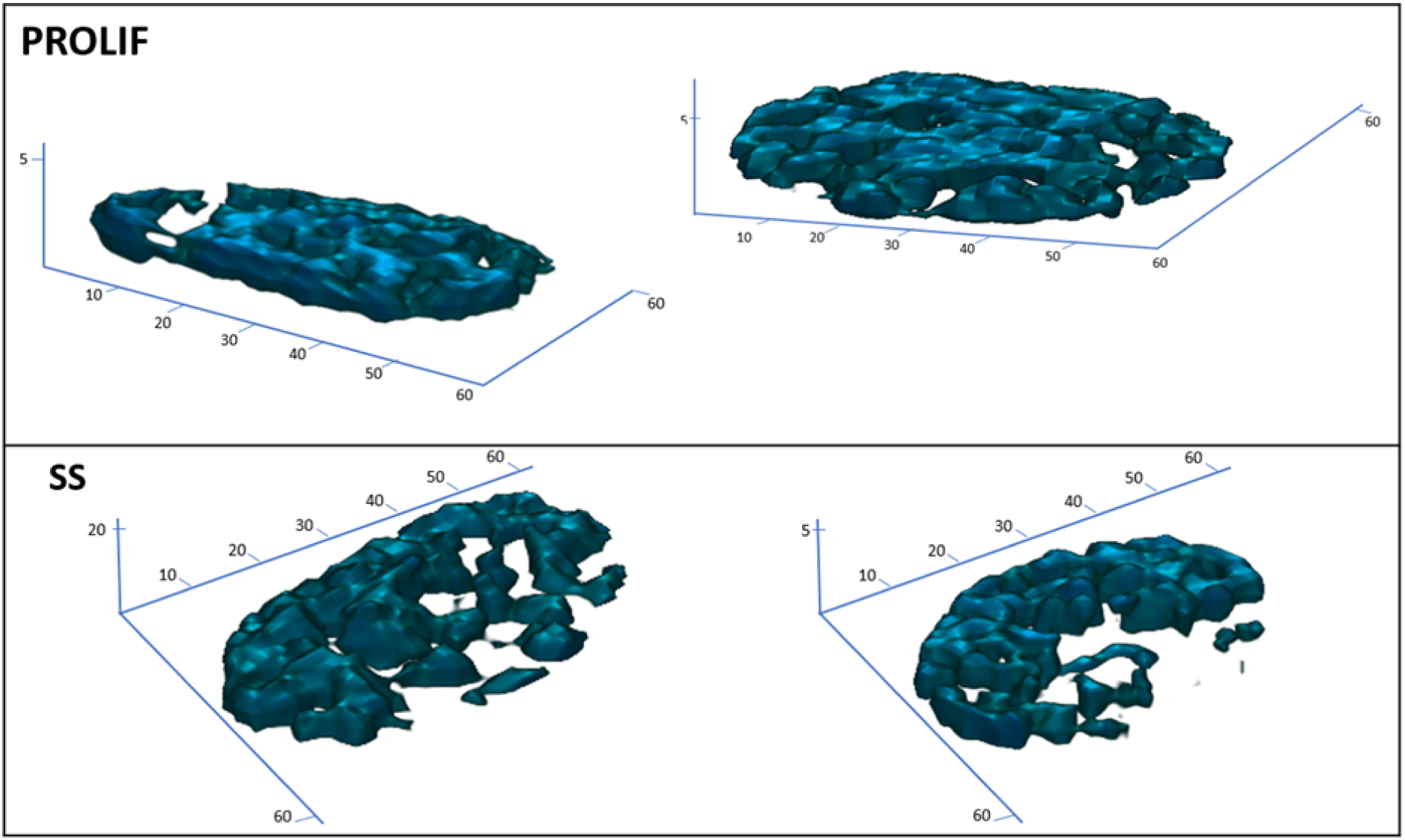
(A) 3D view of Fibroblast Cell Nuclei.

**Fig 8.**
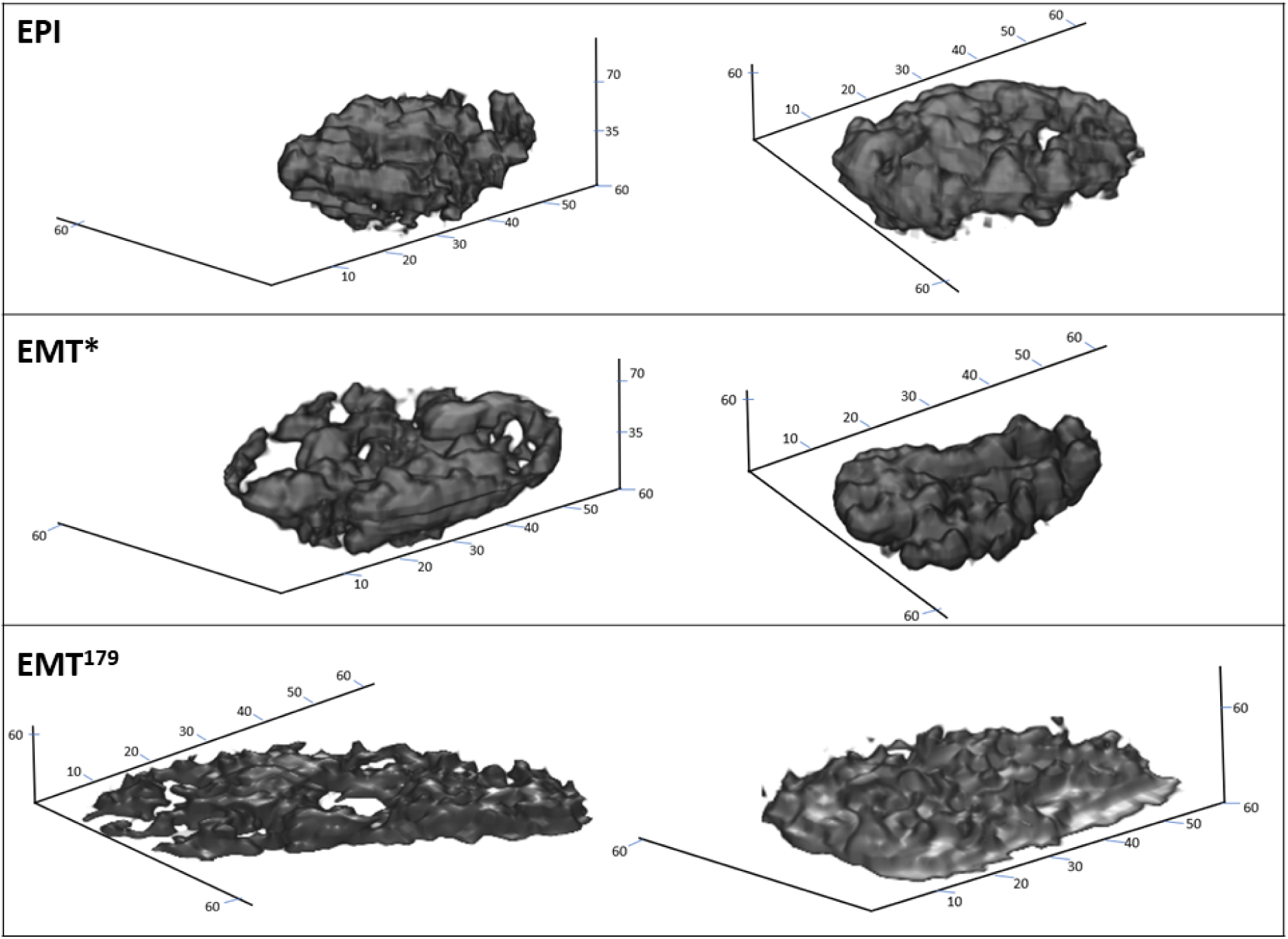
(B) 3D view of PC3 Cell Nuclei.

*EMT* is a mechanism of transition from epithelial to mesenchymal state; it is a reversible process in which case the transition happens from mesenchymal-to-epithelial (MET) as well [38]. During *EMT*, cells do not necessarily exist in ‘pure’ epithelial or mesenchymal state. They comprise of multiple intermediate cell states (*ICS*) that possess hybrid features of both pure states [38]. Therefore, based on obtained results (Table 6), it is suspected that available EMT images consists of cells in two cellular states. While many cells in the *EMT* class of PC3* (*EMT* ^*∗*^) are in the *ICS, EMT* cells in *PC*3^179^ (*EMT* ^179^) are in pure mesenchymal state or in *ICS* close to pure mesenchymal state. Corresponding p-values and z-values characterise cells in *EMT* ^*∗*^ as being of similar intensity but with slightly decondensed HC than in *EPI* state and cells in *EMT* ^179^ as being of lower intensity and highly decondensed HC than in *EPI* and *EMT* ^*∗*^ cells (Fig 8(B)). Previously, the existence of two such *EMT ICS* has been found in mammary epithelium cells [39]. *EMT* cells with their *ICS* are of high importance in studies related to comprehension of cancer progression and drug resistance. Recently Yutong Sha et al. [38] highlighted the characterisation and classification of *ICSs* as a significant challenge and opportunity in the *EMT* field, as they are believed to play a vital role in disease progression and should be considered crucial biological entities. At present single-cell surface marker/RNA sequencing (scRNA-Seq) is the accepted method to identify the intermediate states that occur during *EMT* in metastasis. According to the obtained results, the proposed metrics based on SRP can be utilised to identify and characterise the *ICS* in *EMT* state of human prostate cancer cells. Even though images in *EMT* class consist of cells with diverse characteristics, good classification results are still achieved because of feature representation techniques (BOVW and FV), which derive new image features corresponding to the most dominant original features of the whole data set.

## Conclusion

A 3D nuclear texture description method for cell nucleus classification and variation measurement in chromatin patterns has been presented. This work proposes a new approach to extract 3D texture, where multiple hyperplanes are built within the cubic patch to extract 3D SRP features. The proposed method is utilised to classify nuclear morphology and measure variations in heterochromatin intensity and aggregation (coarsening or opening of heterochromatin) in DAPI images by evaluating the changes in intensity values and differences across two classes. Multiple textural descriptors that range from well known handcrafted texture feature descriptors (SIFT, LBP) to relatively uncommon ones (RSurf, SRP) along with deep learning features (FV-CNN) were also examined. Performance improvement is achieved over the-state-of-the art classification on 3D images of human fibroblast and human prostate cancer cell lines obtained from SOCR, demonstrating the substantial significance of texture features than shape features in the nuclear morphological study. Obtained results also suggest that heterochromatin undergoes considerable change in intensity and aggregation on the transition from normal to another phenotypic state in fibroblast and PC3 cells. Furthermore, two different cellular states in EMT class are identified and characterised, which are considered to hold critical biological significance to study cancer progression and drug resistance.

## Supporting information

**S1 Data**. Available online on the project webpage SOCR 3D Cell Morphometry Project (2018), http://socr.umich.edu/projects/3d-cell-morphometry.

## Acknowledgments

This research includes computations using the computational cluster Katana supported by Research Technology Services at UNSW Sydney.

## Author Contributions

Conceptualisation: Priyanka Rana, Yang Song, Arcot Sowmya, Erik Meijering

Investigation: Priyanka Rana, Yang Song

Methodology: Priyanka Rana, Yang Song

Supervision: Yang Song, Arcot Sowmya, Erik Meijering

Writing - original draft: Priyanka Rana

Writing - review and editing: Priyanka Rana, Yang Song, Erik Meijering, Arcot Sowmya

